# Social risk coding by amygdala activity and connectivity with dorsal anterior cingulate cortex

**DOI:** 10.1101/2024.08.22.608617

**Authors:** Jae-Chang Kim, Leopold Zangemeister, Philippe N. Tobler, Wolfram Schultz, Fabian Grabenhorst

## Abstract

Risk is a fundamental factor affecting individual and social economic decisions, but its neural correlates are largely unexplored in the social domain. The amygdala, together with the dorsal anterior cingulate cortex (dACC), is thought to play a central role in risk taking. Here, we investigated in human volunteers (n=20; 11 females) how risk (defined as variance of reward probability distributions) in a social situation affects decisions and concomitant neural activity as measured with fMRI. We found social variance-risk signals in the amygdala. Activity in lateral parts of the amygdala increased parametrically with social reward variance of the presented options. Behaviorally, 75% of participants were averse to social risk as estimated in a Becker-DeGroot-Marschak auction-like procedure. The stronger this aversion, the more negative was the coupling between risk-related amygdala regions and dACC. This negative relation was significant for social risk attitude but not for the attitude towards variance-risk in juice outcomes. Our results indicate that the amygdala and its coupling with dACC process objective and subjectively evaluated social risk. Moreover, while social risk can be captured with a framework originally established by finance theory for individual risk, the amygdala appears to processes social risk largely separately from individual risk.

## 1. Introduction

Processing risk—the uncertainty associated with outcomes—is crucial for survival and adaptive behavior (Caraco et al., 1980). It allows for an informed prediction of the likely distribution of future outcomes (Knight, 1921). An increase in the spread around the mean of outcomes can serve as an objective measure of an increase in risk (Rothschild and Stiglitz, 1970), operationalized for example as an increase in the variance of outcomes. Decision makers are often not indifferent to increases in objective risk but subjectively evaluate them, e.g. in the form of prospect (Kahneman and Tversky, 1979) and utility (Samuelson, 1937; Houthakker, 1950; Stauffer et al., 2014). The addition of risk reduces the subjective value of a choice option for a risk averse individual but increases it for a risk seeker and these preference differences express themselves in value signals in the brain (Tobler et al., 2009; Van Duijvenvoorde et al., 2015).

Many decisions occur in a social context (Crespi, 2001; Bshary et al., 2014; Chen and Hong, 2018) and the feedback provided by others is valuable to decision makers as it can be a source of acceptance and status (Izuma et al., 2008; Zink et al., 2008). Just like non-social rewards, social rewards are distributed and in principle can be captured with summary statistics such as the mean and variance of the distribution. Accordingly, decision makers may seek or avoid variance-risk in the social domain as such, even without observing the risky choices of others (Chung et al., 2015; Suzuki et al., 2016). Previous research has associated the amygdala and amygdala-dorsal anterior cingulate cortex (dACC) interactions with processing non-social decision variables, including risk (De Martino et al., 2010; Li et al., 2011; Zeeb and Winstanley, 2011; Grabenhorst et al., 2012, 2016; Jung et al., 2013; Orsini et al., 2015, 2017; Chen and Stuphorn, 2018; Feldmanhall et al., 2018; Aydogan et al., 2021). The amygdala processes also social decision variables, such as predictions of social partner’s choices (Grabenhorst et al., 2019), social value (Chang et al., 2015; Schultz et al., 2019), betrayal aversion (Lauharatanahirun et al., 2012) and information about social stimuli and social hierarchy (Gothard et al. 2007; Munuera, Rigotti, and Salzman 2018; Rutishauser et al. 2013). Moreover, it is more active when decision makers align themselves with others in risky decision situations (Burke et al., 2010b). However, the precise role of the amygdala in processing objective or subjectively evaluated social variance-risk remains unclear.

To close this gap, we designed a novel risky choice task in which we first measured the individual willingness-to-pay (WTP) for options with different social and non-social variance-risk as well as different expected value (EV). We used photos together with descriptions of the participant as social rewards and juice as non-social reward, because these two reward types are, or can be made, similar on dimensions such as primacy and duration (Matyjek et al., 2020). For each participant, risk and EV level, we estimated individual level of risk proneness with WTP and its relation to choice. Based on the two separate lines of research described above on amygdala involvement in non-social risk processing and in general social functions, we hypothesized that objective social variance-risk is encoded in the amygdala. Second, we tested whether functional connectivity between amygdala and dACC, suggested by anatomical connections (Carmichael and Price, 1995; Beckmann et al., 2009) and functional similarities (Kable and Glimcher, 2007; McClure et al., 2007; Hare et al., 2011; Kolling et al., 2012), correlated with risk attitude.

## 2. Methods

### 2.1. Participants

After a behavioral session (day1, see below), twenty-four healthy volunteers (age range: 19-29; 14 females) completed the functional magnetic resonance imaging (fMRI) session (day 2) of this study. Inclusion criteria were: normal or corrected-to-normal vision, general liking of dairy products, and normal appetite. Exclusion criteria were: lactose intolerance, active avoidance of sugar or fat in diet, metal implants in body, being on medication other than contraceptives, psychiatric illness, and pregnancy. Four participants were not analyzed further because they showed excessive head motion or because of technical problems during scanning. Accordingly, we present data from 20 participants (11 females). The study was approved by the Local Research Ethics Committee of the Cambridgeshire Health Authority, and written informed consent was obtained from all participants before the experiment.

### 2.2. Tasks

#### 2.2.1. Risky rewards

As social rewards, we used pictures of human faces and the upper half of the upper body with a neutral to positive expression paired with a compliment (Fig. 1a). In our study, social risk consisted of the equiprobable possibility of receiving a larger or a smaller social reward. With higher risk, the magnitude difference between the two rewards was larger than with lower risk. Thus, we used the formal definition of objective risk as the mathematical variance of a known probability distribution (Rothschild and Stiglitz, 1970); for neural applications, e.g., (Tobler et al. 2007, 2009). The subjective values of each social reward were determined in an independent auction-task on Day 1. Individual rewards consisted of three differently flavored commercially available smoothies.

**Figure 1.**
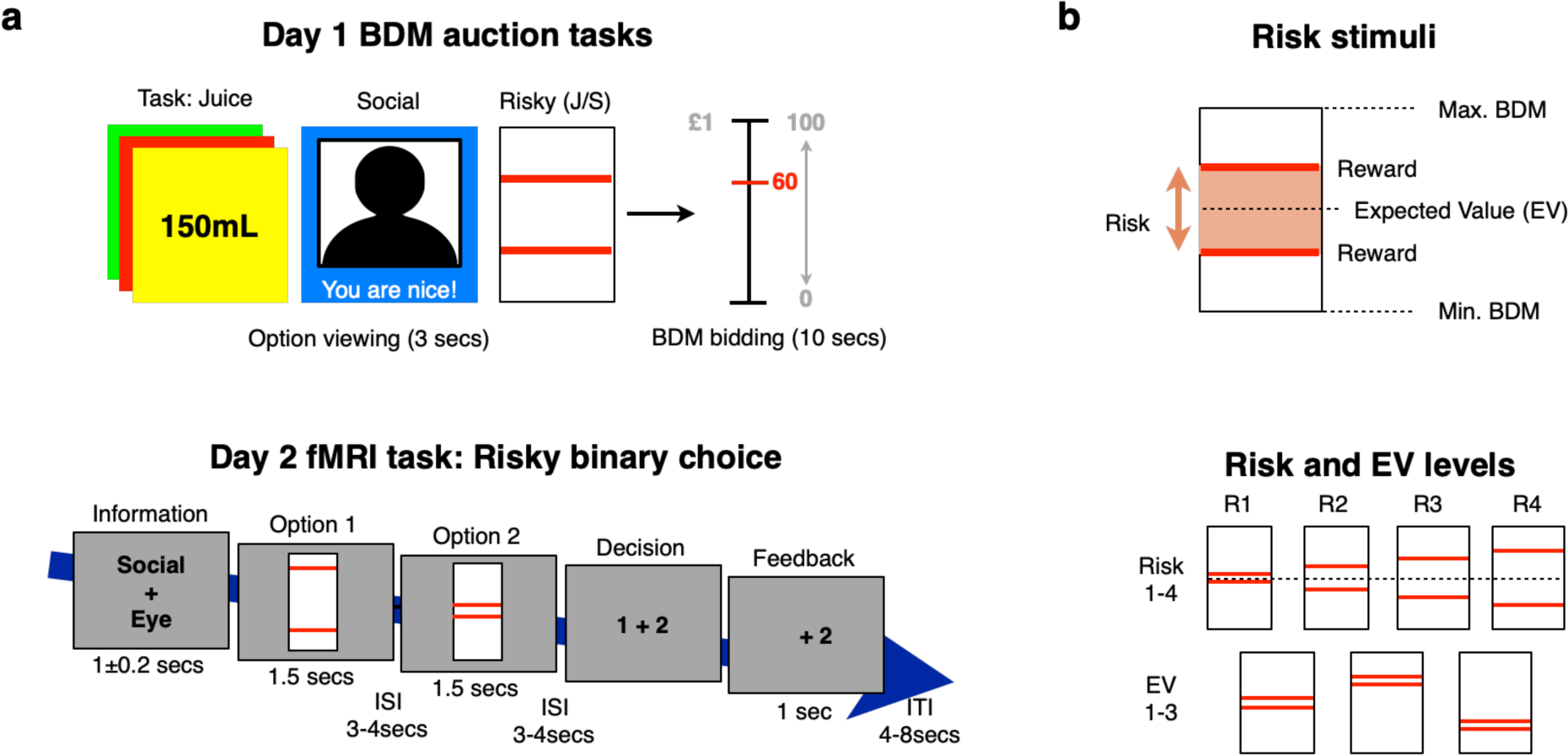
Tasks and stimuli. (a) Tasks. On day 1, all three Becker-De Groot-Marschak (BDM) auction tasks followed the same basic procedure. In each trial, a risk-free (auctions 1 and 2) or a risky option (auction 3) was presented for 3s. Subsequently, participants had up to 10s to indicate their willingness-to-pay on a vertical mouse-controlled scale of 0 to 100 pence in steps of 1 penny (1 penny = £1/100). The possible outcomes illustrated by the bars of risky options corresponded to the amounts participants were willing to pay for specific social or non-social rewards as determined in the first two auction tasks. After each task, one trial was chosen and the participant bid in that trial compared to a computer bid that was randomly drawn from a uniform distribution. If the participant bid in these trials was higher than, or equal to, the computer bid, participants paid the computer bid from their £1 endowment and received the reward at the end of the session. Conversely, if the participant bid was smaller than the computer bid, participants neither paid nor received anything. In the risky choice BDM, participants were informed of the trial type (social / juice) before being presented with the risky option. Note that actual photographs were used for social trials in the study. On day 2, participants performed the fMRI task, in which they chose between two risky options on every trial, either by eye movement (saccade) or by hand movement (button press). We used different actions to keep participants engaged. Both reward type (juice or social) and action type varied across trials. The trial type information (here “Social + Eye”; presented for 1s ±200ms) was followed by the sequential presentation of the two options (presented for 1.5s each). Participants then provided their choice (option 1 represented by “1” and option 2 by “2”) with the trial-appropriate action. In the feedback phase, the chosen option remained visible for 1s. Jittered intervals separated trials by 4-8s and stimuli by 3-4s. (b) Stimuli. Our stimuli consisted of a rectangular box with two horizontal bars, each representing a reward. These two rewards were equiprobable (p=0.5). The risky options in the main task were the same as those used in the third BDM auction. Risk levels R1-4 reflect different levels of variance-risk. These were crossed with three levels of expected value, EV1-3, defined as the mean of the two equiprobable rewards. Thus, overall, there were 12 different options (3 EV levels times 4 risk levels), both for the social and the non-social domain. Abbreviations: EV, expected value; ISI, inter-stimulus-interval; ITI, inter-trial-interval.

In the auction task on Day 1 and the choice task on Day 2, we presented social and individual options with a specific risk and EV. As usual, the expected value of a gamble corresponded to the sum of the probability-weighted reward amounts that could be won in that gamble (Knutson et al., 2005; Tobler et al., 2005, 2009). Because our gambles all consisted of two equiprobable rewards, the expected value of a gamble was simply the mean of the two possible reward amounts. We used four levels of risk and three levels of EV, resulting in 12 unique risk-EV combinations. Each option consisted of a rectangular box and two red horizontal lines at different heights (Fig. 1b), corresponding to different reward amounts (O’Neill and Schultz, 2010). Participants learned the social and individual reward levels associated with line height on Day 1. The same stimuli informed participants about social and individual risk in the fMRI task on Day 2. Therefore, differential brain activity to social and individual risk cannot be due to visual differences in the stimuli.

#### 2.2.2. Becker-DeGroot-Marschak (BDM) auction-like task (Day 1)

On day 1, participants evaluated social and non-social (juice) rewards in a social and a juice valuation task. Specifically, they used a computer mouse to provide a bid (b) on how much money they were willing to pay in order to obtain a given social or juice reward. Before each reward task, participants received an endowment of £1. Each trial consisted of a reward presentation phase of 3s followed by a bidding phase of up to 10s. Rewards were kept constant across participants. Juice rewards were cued by a rectangle in one of three different colors, indicating the type of juice (Matyjek et al., 2020), and an amount, indicated in ml. These amounts ranged from 15 to 330ml in steps of 15ml.

For the valuation of social rewards, each face and each compliment appeared only once. For each reward type, there were 66 unique rewards, yielding 66 unique reward-willingness-to-pay combinations. Following a standard Becker-DeGroot-Marschak (BDM) auction (Becker et al., 1964), the actual price of the reward in one randomly selected trial was determined by a uniformly distributed random draw from 0 to 100 pence (1 penny = £1/100). If the bid of the participant was larger than, or equal to the actual price, the participant received the reward and paid the actual price. In contrast, if the bid of the participant was smaller than the price, the participant did not receive the reward but did not pay anything either. One trial of each auction task was implemented at the end of the session.

Next, to measure the subjective value of variance-risk, participants provided bids to play various risky options with two equiprobable rewards in a third BDM auction task. Before each trial, participants were notified about the reward type (social or juice) they were bidding for in that trial. Participants evaluated all twelve options, both for social and juice reward. Again, the computer randomly selected one trial at the end of the task and this trial was implemented according to the randomly determined actual price and the bid of the participant. Finally, participants practiced the main task to be used on Day 2 (see next section).

#### 2.2.3. fMRI task: Risky binary choice (Day 2)

On Day 2, participants first reacquainted themselves with the main task inside the MRI scanner. Participants performed three blocks of 48 trials (17.73±0.34 minutes per block, mean±SEM). In every trial of this task, participants decided between two risky options (Fig. 1a). These options were the same as those used in the risky gamble BDM task on Day 1. Each trial started with information (1±0.2 s) about its reward (social or juice) and required action type (eye or hand movement). Subjects then saw two consecutively presented risky options (1.5 s each). The two options were separated by a jittered interstimulus interval (fixation cross, 3-4 s). All choices were between a low risk (level 1 in Fig. 1b) and a higher risk (levels 2-4) option, presented in randomized order, with an equal number of trials in which the low risk option appeared first or second. The EV of the low risk option varied from trial-to-trial, which prevented participants from fully predicting the properties of the second option even if it was low risk and thereby precluded decision-making already during the presentation of the first option. The sequential presentation of options allowed us to look for neural correlates of single-option risk and EV. After viewing the options, participants chose either the option they saw first (“1”) or the option they saw second (“2”). The “1” and the “2” were randomly presented to the left and right of a fixation cross, preventing movement preparation already at the time of the second option. The placeholder of the chosen option (“1” or “2”) remained on the screen for 1 s after the choice. Trials were separated by jittered inter-trial-intervals (ITI) of 4-8 s. Both ISIs and ITIs were jittered according to a Poisson distribution to increase design efficiency.

To keep participants engaged, we asked them to provide their choices either by button press on button box or by saccade. Using the word “eye” or a “hand”, participants were informed of the required action type at the beginning of each trial, together with the juice/social information. There was a fixation requirement in all trials such that if participants made a saccade in a hand trial or a button press in an eye trial, the trial was counted as error and repeated. Before the MRI task, we informed participants that one randomly selected decision per reward category would be played out with the risk and EV of the selected option. Participants received the rewards they won after the MRI session in the form of a glass of chosen juice reward and a printed card with the social reward.

### 2.3. Neuroimaging

#### 2.3.1. Data acquisition

Participants were scanned with a Siemens 3T Trio Scanner at the Cognition and Brain Sciences Unit, Cambridge, UK. We acquired between 310 and 420 volumes of T2*-weighted echo-planar images (EPIs) in three runs. For each volume, we acquired 56 slices in an ascending order and with the following parameters: in plane resolution 3 × 3 mm, slice thickness 2 mm, repetition time (TR) = 3 s, echo time (TE) = 30 ms, flip angle = 90°, slice gap = 0.5mm, field of view (FOV) = 192 x 192 x 140mm^3^, matrix = 64 x 64. Four dummy volumes before each scanning run served to achieve steady-state magnetization. We acquired a high-resolution T1 structural scans using a MPRAGE sequence, 192 slices, slice thickness 1mm, no gap, in-plane resolution, 1 x 1 mm^2^, FOV = 256 x 256 x 192 mm^3^, TR = 2.3 s, TE = 2.98 ms, inversion time 900 ms, flip angle = 9°.

#### 2.3.2. Image preprocessing

We used Statistical Parametric Mapping (SPM12) to preprocess the fMRI data, including slice-time correction and motion correction. All pre-processed fMRI data were aligned to the high-resolution anatomical image. After segmentation of the anatomical images into gray and white matter, they and the co-registered EPI images were DARTEL-normalized to MNI-space. Finally, the EPI images were spatially smoothed using a three-dimensional Gaussian filter (6 mm full-width-at-half-maximum).

### 2.4. Behavioral data analysis

#### 2.4.1. BDM auction tasks

To assess the influence of objective variance-risk, expected value and their interaction on individual willingness-to-pay (i.e., subjective value), we performed the following multiple linear regression for each participant, separately for the social and non-social domain:

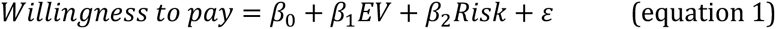

where 𝛽_0_ refers to the intercept, 𝛽_1_ and 𝛽_2_ refer to the model-specific fixed effect regression coefficients (beta weights), and 𝜀 refers to the residual error. Participants with 𝛽_2_ < 0 are risk averse, participants with 𝛽_2_ > 0 risk seeking.

#### 2.4.2. Risky binary choice task

To assess the influence of the willingness to pay (derived from the separate risky option BDM auction task; WTP) for each option, we performed the following logistic regression:

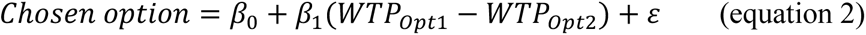

With the chosen option as either 0 or 1 (chose option 1 or option 2, respectively) 𝑊𝑇𝑃*_opt_*_1_ − 𝑊𝑇𝑃*_opt_*_2_ as the signed difference between the expected values of option 1 and option 2, 𝛽_0_ as the constant term, 𝛽_1_ as the corresponding slope parameter estimate, and 𝜀 as the residual.

#### 2.4.3. Risk proneness and EV leaning across tasks

To quantify individual proneness to take social or non-social risk, we used the regression coefficients 𝛽_2_ from the equation (1) above (section 2.4.1) for each participant. For completeness, we also considered the regression coefficients 𝛽_1_(equation (1) in section 2.4.1) to quantify individual sensitivity to social and non-social EV.

### 2.5. Neuroimaging data analysis

To detect neural activity related to variance-risk, EV as well as to WTP, we estimated two general linear models (GLM) in SPM12 at the participant (first) level and then took individual contrast images to the group (second) level.

#### 2.5.1. Participant level analysis (1^st^ level)

In the first GLM (GLM1), for each participant and block of trials, we modelled five main time periods within each trial: 1) information screen onset (juice/social reward, eye/hand action); 2) social option 1 onset; 3) social option 2 onset; 4) non-social option 1 onset; 5) non-social option 2 onset; 6) time of decision; and 7) feedback screen onset. Periods 2-5 each had two parametric modulators, one for social or non-social variance-risk and the other for social or non-social expected value. To ensure that the regressors competed for explaining independent components of variance, serial orthogonalization of parametric regressors was turned off (Mumford et al., 2015). We modelled each period as an event with a duration of zero. Each of these regressors was convolved with the canonical basis functions. Furthermore, we added the six unconvolved motion regressors and a regressor for each block to account for potential confounds. Simple contrasts on parametric modulators for social or non-social risk (averaged across option 1 and 2) served to construct individual contrast images.

In the second GLM (GLM2), we modelled both social and non-social as one trial type, and included a parametric modulator, reflecting WTP as determined via a Becker-DeGroot-Marschak (BDM) auction-like task. We conducted the following multiple linear regression for each participant, analyzing the domain-general WTP coding. Similar to GLM1 above, we modelled each period as an event (duration of zero), convolving all regressors with canonical basis functions. Motion regressors and a block regressor were added to address potential confounds. Individual contrast images were constructed using simple contrasts on parametric modulators for WTP.

#### 2.5.2. Group analysis (2^nd^ level)

To identify regions processing social or non-social variance-risk at the group level, we used flexible factorial designs that included the contrast images on the parametric modulators from the individual GLM. We report whole-brain-results (p < 0.05, voxel-level FWE-corrected) as well as activations in the amygdala, our a priori region of interest (p < 0.05, peak-level FWE corrected). To define the region of interest, we used the third version of the automated anatomical labelling atlas (AAL 3; Rolls et al. 2020). To extract activity from amygdala regions we used right amygdala from AAL3.

#### 2.5.3. Functional connectivity between amygdala and medial prefrontal cortex

To assess risk level-dependent functional interactions between the amygdala and medial prefrontal cortex during risky decision making, we performed a psychophysiological interaction (PPI) analysis (Friston et al., 1997; Gitelman et al., 2003). For each participant, we first extracted the eigenvariate time series from the right amygdala as seed region. The signals were de-convolved to construct a time series capturing neural activity in the seed area, which served as the physiological regressor for the PPI analyses. As psychological regressors, we used 1) social variance-risk, 2) juice variance-risk. The interaction regressors corresponded to the multiplication of the physiological and psychological regressors. Again, we added the six motion regressors and block regressors of no interest. Next, we correlated the social or juice risk-related interaction contrast images with the individual proneness for social or juice risk (𝛽_2_ of the equation (1) in section 2.4.1). For this analysis, we focused on connectivity of the amygdala with dorsal anterior cingulate cortex (dACC), because this connection appears to be particularly important for risk taking (Jung et al., 2018). The dACC ROI was defined with the Connectivity-based atlas (Neubert et al., 2015) particularly anterior rostral cingulate zone. We also defined a ROI for the temporoparietal junction (TPJ) area (Mars et al., 2012), because of previous literature implicating this region in aspects of social cognition (Hill et al., 2019).

## 3. Results

To investigate whether and how the amygdala, in possible interaction with the cortical regions, processes social variance-risk, we asked participants to choose between two sequentially presented options with different variance-risk and expected value. We had determined the subjective value of the available social and non-social rewards, as well as the willingness of participants to take social and non-social variance-risk, in separate bidding tasks.

### 3.1. Behavioral Results

Our bidding tasks (Becker-DeGroot-Marschak auctions) indicated that our participants were willing to pay for social or juice rewards and showed considerable individual variation in their valuation of variance-risk (equation (1) in section 2.4.1), both in the social and the non-social domain (Fig. 2). When we averaged risk attitudes across social and non-social domains (Fig. 2a), 12 out of our 20 participants were risk averse (i. e., 𝛽_2_ (from equation 1) < 0, decreasing willingness to pay with higher variance-risk). The remaining 8 were risk seeking (𝛽_2_ (from equation 1) > 0, increasing willingness to pay with higher variance-risk). Thus, the majority of participants was risk averse, although there was also a sizeable number of risk seeking participants. A similar picture emerged when we analyzed risk attitudes separately for the social (Fig. 2b) and non-social (Fig. 2c) domain. Fifteen participants were averse to social variance-risk and 11 to juice variance-risk. Risk attitudes in the two domains were correlated (Spearman’s rho =0.52, p=0.02). These data suggest that participants processed variance-risk and that they did so according to individual preferences, not only when this risk concerned juice reward but also when it concerned social reward and that risk attitude correlated between social and non-social risk.

**Figure 2.**
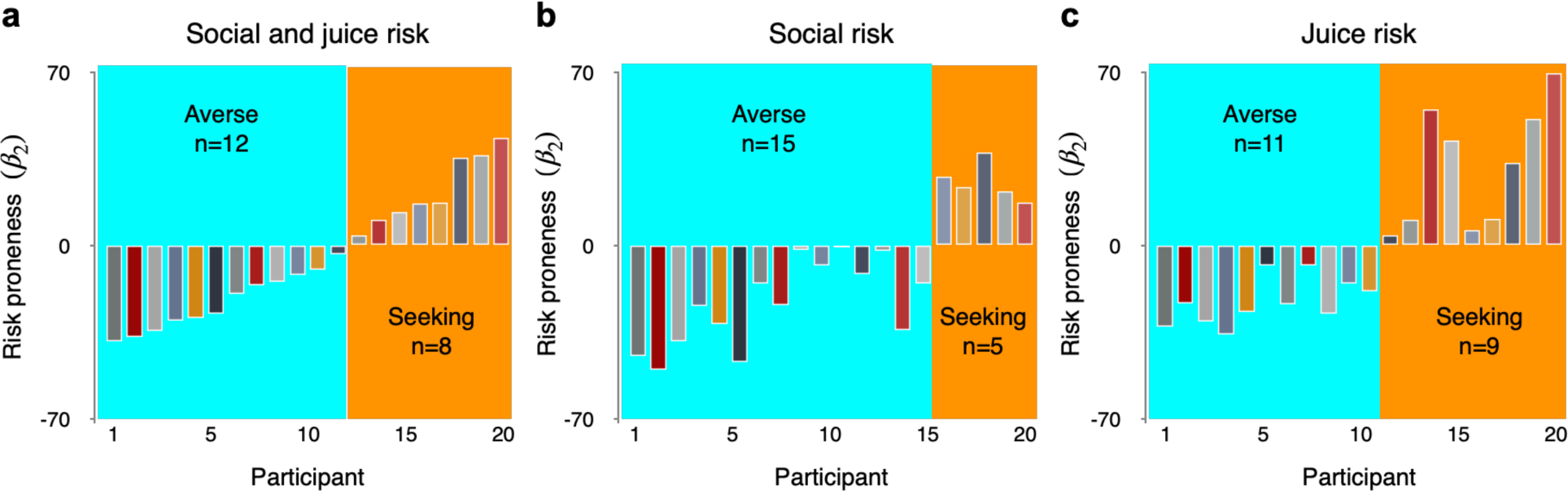
Risk proneness across and within domains. (a) Risk proneness averaged across social and non-social domains. Twelve participants were risk averse (𝛽_2_< 0; light blue), eight risk seeking (𝛽_2_> 0; orange). Participants are ordered from least risk-prone to most risk-prone. (b) Risk proneness in the social domain. (c) Risk proneness for juice variance-risk. In (b) and (c), the ordering of participants was the same as in (a).

In a supplementary analysis, we tested whether willingness-to-pay was a predictor of binary choices (equation 2 in section 2.4.2). Indeed, we found that WTP for risky options predicted binary choices between risky options (logistic regression; group-level t-test; P = 0.05, t(19) = 2.08). A more detailed analysis of single participant regression coefficients revealed that this was only the case for 15 out of 20 participants. This finding suggested that WTP bids for risky options can predict binary choices, but that the congruence does not occur in all participants. The lack of congruence in these participants was likely due to additional value comparisons between binary choice options beyond the simple WTP assigned to individual options. Nevertheless, we included all 20 participants for the fMRI analyses of risk, expected value, and WTP.

### 3.2. Neural Results

#### 3.2.1. Social variance-risk coding in amygdala

To test whether amygdala activity correlated with social variance-risk, we regressed blood oxygen level-dependent (BOLD) responses against trial-by-trial parametric modulators capturing variance-risk when the two options were presented (‘neuroimaging data analysis’ in Methods). Amygdala activity increased with variance-risk for both social (small-volume FWE peak-corrected, p<0.05; peak Montreal Neurological Institute (MNI) coordinate X=-18,Y=-4,Z=-12 for left amygdala: t=4.97; X=32,Y=0,Z=-16 for right amygdala: t=4.01; Fig. 3a, yellow) and juice reward (small-volume FWE peak-corrected, p<0.05; X=24,Y=2,Z=-22 for right amygdala: t=4.18; Fig. 3a, green). The direct comparisons between social and juice risk showed no significant effects after correction. Furthermore, to assess common activations related to both social and non-social risk, we used conjunction analysis and found domain-general risk signals in the amygdala, as well as other regions (e.g., OFC; Table 2). The correlation with aversion to social risk at this location was not significant (t=1.16, p=0.5). Moreover, we found no relation to risk proneness or risk aversion in any other voxel of the amygdala, even at lenient thresholds (p<0.005, uncorrected). Thus, the amygdala processed social and juice variance-risk.

**Figure 3.**
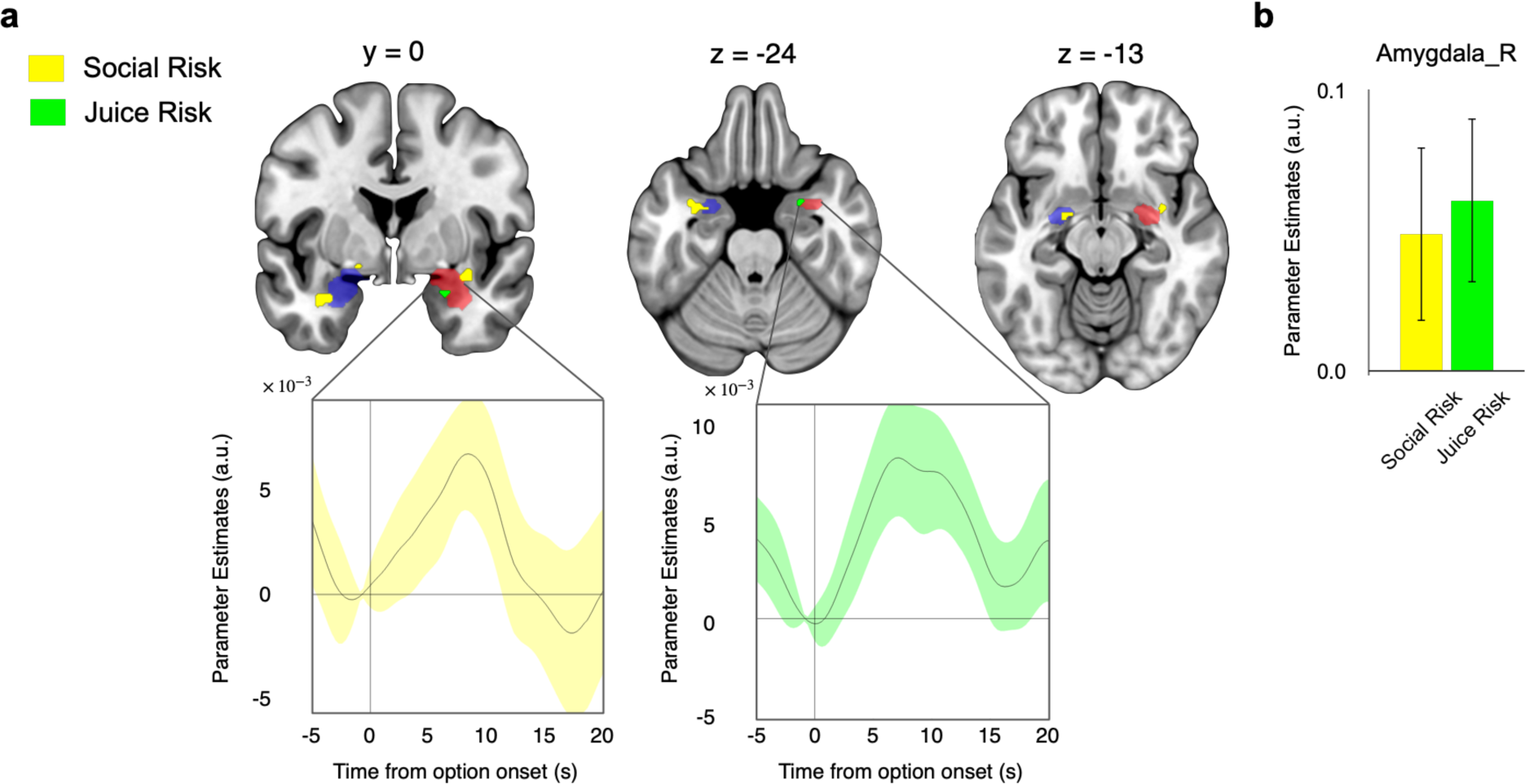
Variance-risk signals in amygdala. (a) Location of variance-risk signals. Social variance-risk signals (left amygdala: t=4.97; right amygdala: t=4.01; yellow) located bilaterally and more laterally whereas juice variance-risk signals (right amygdala: t=4.18; green) were unilateral and located more medially. Neural activations are small volume FWE peak-level corrected p<0.05). (b) Parameter estimates for entire right amygdala (red in (a)). Responses to the risky options increased with both social and juice variance-risk. Error bars represent ±1 standard error of the mean.

**Table 1.** Risk proneness 𝛽_2_ and EV sensitivity 𝛽_1_ across and within domains.

| Participant<br># | Social and juice<br>Risk proneness ($\beta_2$) | $p$ | Social and juice<br>EV sensitivity ($\beta_1$) | $p$ | $R^2$ |
| --- | --- | --- | --- | --- | --- |
| 1 | -24.36 | 0.000 | -26.03 | 0.445 | 0.36 |
| 2 | -22.35 | 0.000 | 116.13 | 0.000 | 0.70 |
| 3 | -14.21 | 0.000 | 115.75 | 0.000 | 0.53 |
| 4 | -11.94 | 0.016 | 130.88 | 0.000 | 0.33 |
| 5 | -11.52 | 0.000 | 84.28 | 0.000 | 0.47 |
| 6 | -8.21 | 0.001 | 72.38 | 0.000 | 0.40 |
| 7 | -5.61 | 0.087 | 86.13 | 0.000 | 0.29 |
| 8 | -5.29 | 0.010 | 74.06 | 0.000 | 0.45 |
| 9 | -3.74 | 0.000 | 100.22 | 0.000 | 0.87 |
| 10 | -3.06 | 0.003 | 178.56 | 0.000 | 0.94 |
| 11 | -2.68 | 0.647 | 28.56 | 0.482 | 0.02 |
| 12 | -0.06 | 0.979 | 175.66 | 0.000 | 0.76 |
| 13 | 0.43 | 0.689 | 95.56 | 0.000 | 0.79 |
| 14 | 0.79 | 0.909 | 90.28 | 0.066 | 0.07 |
| 15 | 1.70 | 0.302 | 69.56 | 0.000 | 0.46 |
| 16 | 3.12 | 0.113 | 168.53 | 0.000 | 0.78 |
| 17 | 3.51 | 0.013 | 47.94 | 0.000 | 0.42 |
| 18 | 11.92 | 0.000 | 74.69 | 0.000 | 0.73 |
| 19 | 15.57 | 0.000 | 35.75 | 0.033 | 0.52 |
| 20 | 22.28 | 0.000 | 13.38 | 0.650 | 0.39 |

| Participant<br># | Social<br>Risk proneness ($\beta_2$) | $p$ | Social<br>EV sensitivity ($\beta_1$) | $p$ | $R^2$ |
| --- | --- | --- | --- | --- | --- |
| 1 | -20.95 | 0.003 | -31.63 | 0.481 | 0.35 |
| 2 | -21.63 | 0.000 | 129.88 | 0.000 | 0.83 |
| 3 | -15.21 | 0.002 | 127.69 | 0.000 | 0.59 |
| 4 | -15.58 | 0.028 | 149.94 | 0.004 | 0.43 |
| 5 | -10.14 | 0.023 | 71.75 | 0.021 | 0.37 |
| 6 | -8.39 | 0.001 | 68.63 | 0.000 | 0.62 |
| 7 | -5.97 | 0.062 | 50.94 | 0.025 | 0.32 |
| 8 | -7.88 | 0.012 | 63.50 | 0.004 | 0.46 |
| 9 | -3.80 | 0.003 | 103.06 | 0.000 | 0.89 |
| 10 | -4.05 | 0.019 | 164.50 | 0.000 | 0.92 |
| 11 | -9.48 | 0.002 | 66.56 | 0.002 | 0.54 |
| 12 | -1.80 | 0.508 | 170.69 | 0.000 | 0.80 |
| 13 | -1.00 | 0.558 | 89.38 | 0.000 | 0.74 |
| 14 | 3.36 | 0.236 | 49.44 | 0.018 | 0.28 |
| 15 | 5.36 | 0.020 | 79.00 | 0.000 | 0.63 |
| 16 | 1.46 | 0.573 | 159.69 | 0.000 | 0.80 |
| 17 | 5.11 | 0.001 | 35.00 | 0.001 | 0.61 |

| 18 | 14.01 | 0.000 | 92.81 | 0.000 | 0.75 |
| --- | --- | --- | --- | --- | --- |
| 19 | 11.23 | 0.000 | 44.75 | 0.014 | 0.58 |
| 20 | 10.25 | 0.063 | -12.00 | 0.745 | 0.16 |
| Participant # | Juice Risk proneness ($\beta_2$) | $p$ | Juice EV sensitivity ($\beta_1$) | $p$ | $R^2$ |
| 1 | -27.78 | 0.000 | -20.44 | 0.620 | 0.52 |
| 2 | -23.07 | 0.000 | 102.38 | 0.001 | 0.72 |
| 3 | -13.21 | 0.010 | 103.81 | 0.004 | 0.47 |
| 4 | -8.29 | 0.234 | 111.81 | 0.027 | 0.25 |
| 5 | -12.91 | 0.000 | 96.81 | 0.000 | 0.71 |
| 6 | -8.03 | 0.044 | 76.13 | 0.008 | 0.39 |
| 7 | -5.24 | 0.226 | 121.31 | 0.001 | 0.47 |
| 8 | -2.69 | 0.300 | 84.63 | 0.000 | 0.53 |
| 9 | -3.68 | 0.014 | 97.38 | 0.000 | 0.84 |
| 10 | -2.07 | 0.046 | 192.63 | 0.000 | 0.97 |
| 11 | 4.13 | 0.435 | -9.44 | 0.797 | 0.03 |
| 12 | 1.69 | 0.549 | 180.63 | 0.000 | 0.81 |
| 13 | 1.86 | 0.165 | 101.75 | 0.000 | 0.86 |
| 14 | -1.78 | 0.424 | 131.13 | 0.000 | 0.78 |
| 15 | -1.97 | 0.147 | 60.13 | 0.000 | 0.69 |
| 16 | 4.78 | 0.127 | 177.38 | 0.000 | 0.78 |
| 17 | 1.91 | 0.357 | 60.88 | 0.000 | 0.48 |
| 18 | 9.82 | 0.000 | 56.56 | 0.000 | 0.80 |
| 19 | 19.90 | 0.000 | 26.75 | 0.168 | 0.73 |
| 20 | 34.30 | 0.000 | 38.75 | 0.194 | 0.77 |

Participant # is sorted by low risk proneness to high risk proneness across domains.

**Table 2.**
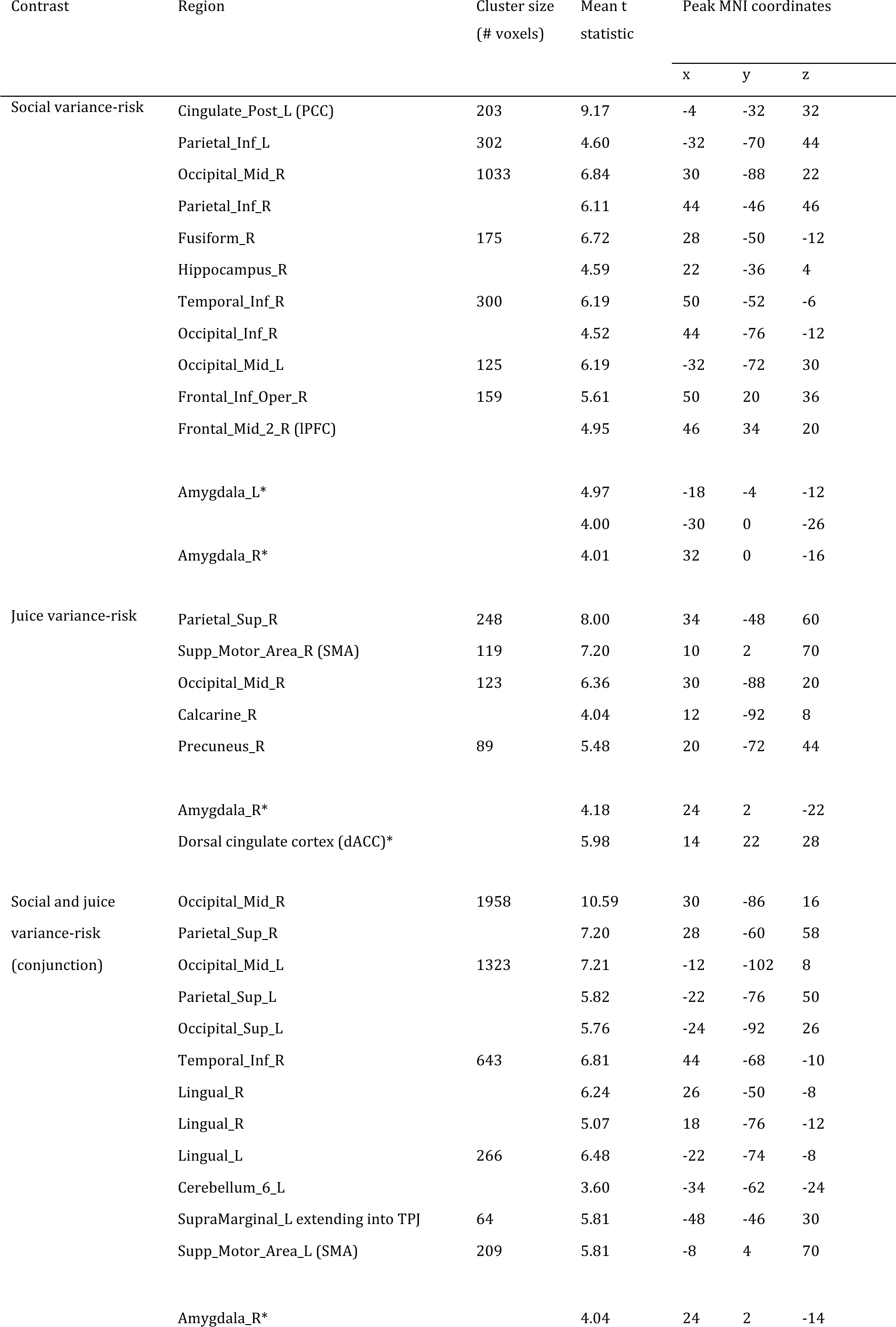
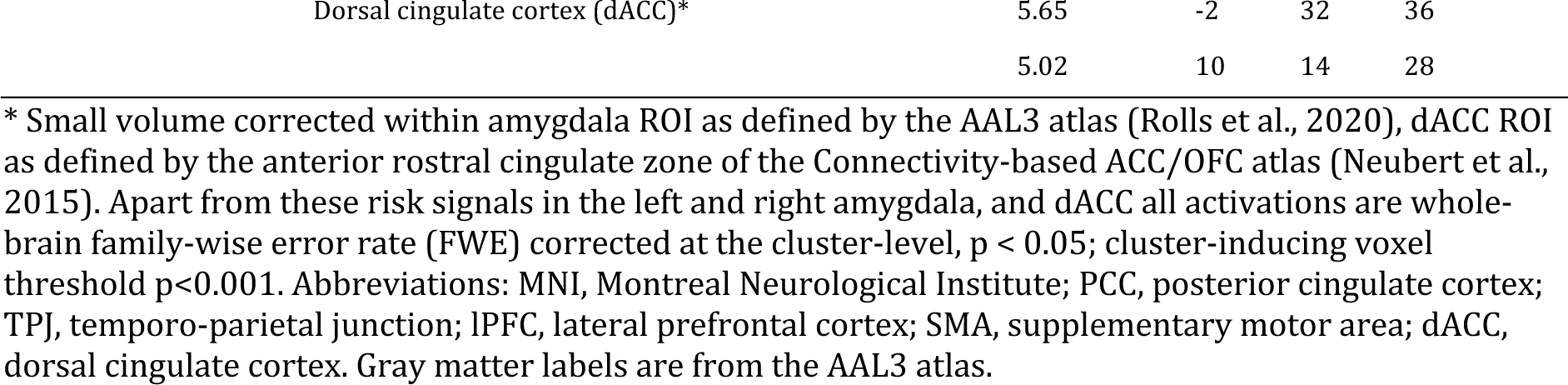
Brain regions processing social or non-social variance-risk.

In exploratory analyses, we investigated social and non-social variance risk coding in the anterior insular cortex, striatum, and lateral orbitofrontal cortex (OFC), as these regions have been associated with risk processing previously (Preuschoff et al., 2006; Tobler et al., 2007; Cooper and Knutson, 2008; Burke et al., 2010b; O’Neill and Schultz, 2010, 2013). We observed significant social variance risk signals in the striatum (small-volume FWE-corrected, p<0.05), specifically in nucleus accumbens and putamen, as well as in the OFC (small-volume FWE-corrected, p<0.05 in combination of medial, anterior, posterior, and lateral OFC). The insular cortex showed a relation to social risk that did not survive correction (small-volume FWE-corrected, p=0.088). Relations to non-social risk occurred in the striatum (small-volume FWE-corrected, p<0.05), and did not reach significance in insula and OFC (both p<0.005 uncorrected).

#### 3.2.2. WTP coding in mPFC, middle and posterior cingulate cortex

We found whole-brain FWE cluster-corrected WTP signals (Fig. 4) in the medial prefrontal cortex (mPFC; t=6.87), middle cingulate cortex (MCC; t=7.44) and posterior cingulate cortex (PCC; t=6.25). Other regions showing WTP signals included the occipital regions (Occipital_Inf_R, t=7.85; Lingual_R, t=6.69; Cuneus_L, t=7.55) and SMA (t=6.38; Table 3). Furthermore, we found WTP signals in the right amygdala (small-volume FWE peak-corrected, p<0.05; X=24,Y=-4,Z=-18 for right amygdala). Thus, consistent with frequently identified value-processing regions (Bartra et al., 2013), we found WTP signals in parts of the medial prefrontal and cingulate cortex.

**Figure 4.**
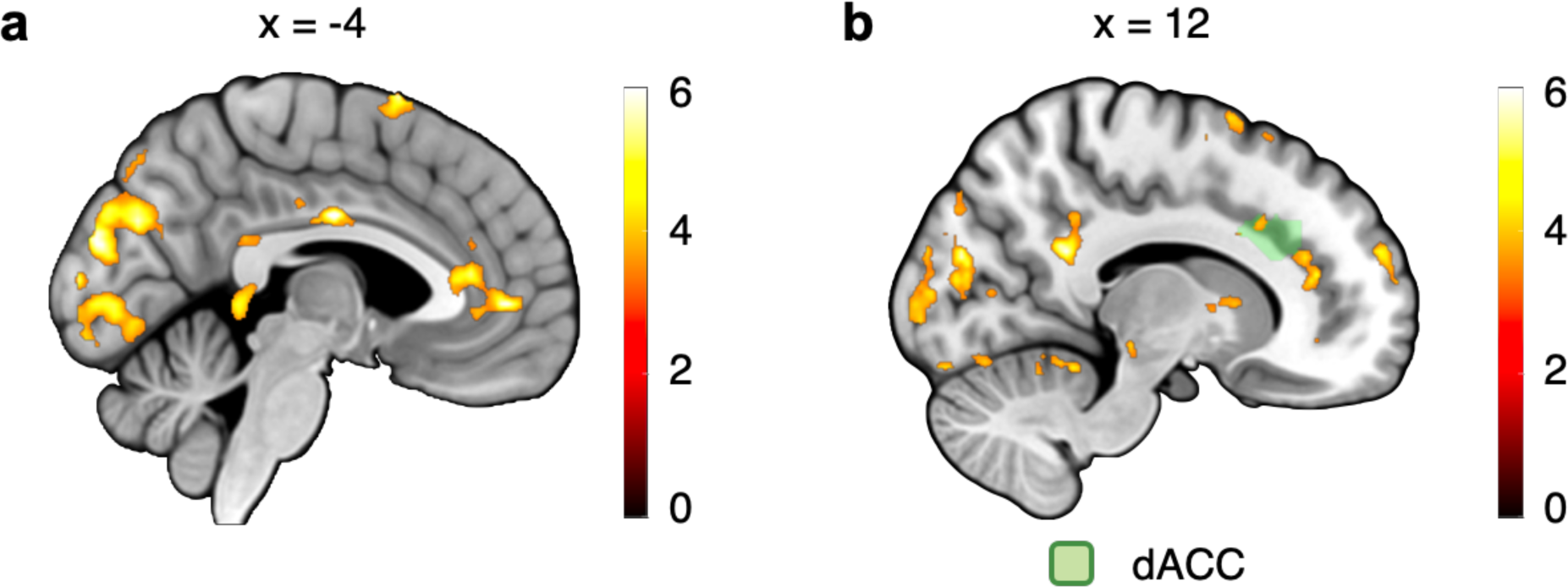
Neural representation of willingness-to-pay (WTP). (a) WTP parametrically activated MCC (t=7.44), PCC (t=6.25), mPFC (t=6.87), SMA (t=6.38), and occipital cortex (Occipital_Inf_R, t=7.85; Lingual_R, t=6.69; Cuneus_L, t=7.55). Neural activations are whole-brain FWE cluster-corrected p<0.05, cluster-inducing voxel threshold p<0.001. (b) The dorsal anterior cingulate cortex (dACC, t=4.50) also encodes WTP. Neural activations are small volume FWE peak-level corrected with p<0.05 (cluster-forming threshold: p<0.001).

**Table 3.**
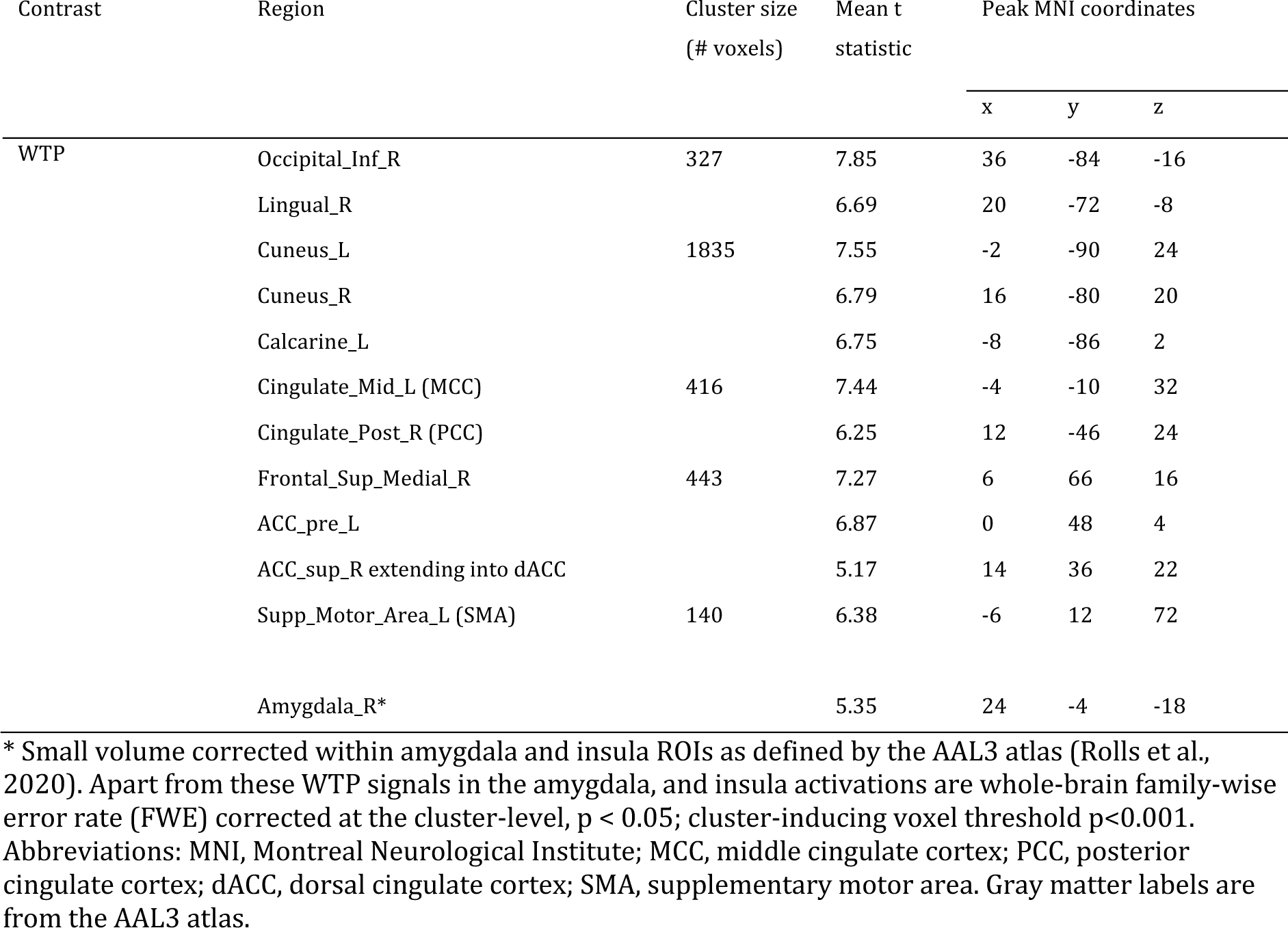
Brain regions processing WTP.

#### 3.2.3. Negative functional coupling between amygdala-dACC, -TPJ and subjective risk proneness

Our activation results relate the right amygdala to variance-risk for both juice and social reward. Previous studies have shown that the amygdala is anatomically (Carmichael and Price, 1995; Beckmann et al., 2009) and functionally (e.g., (Apps et al., 2016) connected to the dorsal anterior cingulate cortex; dACC). Moreover, resting-state connectivity between the amygdala to the other regions (i.e., node strength) and particularly dACC seems to be related to attitudes concerning non-social risk positively (Jung et al., 2018). However, the relation of task-induced activity to taking either social or non-social risk remains unknown. We therefore investigated whether amygdala-dACC connectivity during the presentation of risky choice options changed as a function of risk proneness.

In our psychophysiological interaction (PPI) connectivity analysis, we used right amygdala activity as physiological regressor and parametric variance-risk as psychological regressor. This analysis revealed that a lower proneness to take social variance-risk correlated with stronger connectivity to the dACC (p<0.05, peak-level FWE small-volume corrected within the dACC as defined by the anterior rostral cingulate zone of the Connectivity-based ACC/OFC atlas (Neubert et al., 2015), with a cluster-forming threshold of p<0.001).

We also observed negative functional coupling between amygdala-TPJ area and social risk proneness regression coefficients 𝛽_2_ from the equation above (section 2.4.1) for each participant. The coupling in right TPJ withstood whole-brain FWE cluster-level correction (p < 0.05, cluster-forming threshold: p<0.001; Fig. 5). Additionally, the variance-risk associated with juice did not persist in either dACC or TPJ when thresholded at p<0.001. The entire right amygdala was chosen as a seed region due to increased activity observed in response to both social and juice variance-risk (Fig. 3). We used the entire right amygdala as a seed region because activity in that area increased for both social and juice variance-risk (Fig. 3). These findings suggest that risk proneness correlates with negative coupling between the amygdala and both dACC and TPJ when processing social risk, but this is not the case with risk for juice reward.

**Figure 5.**
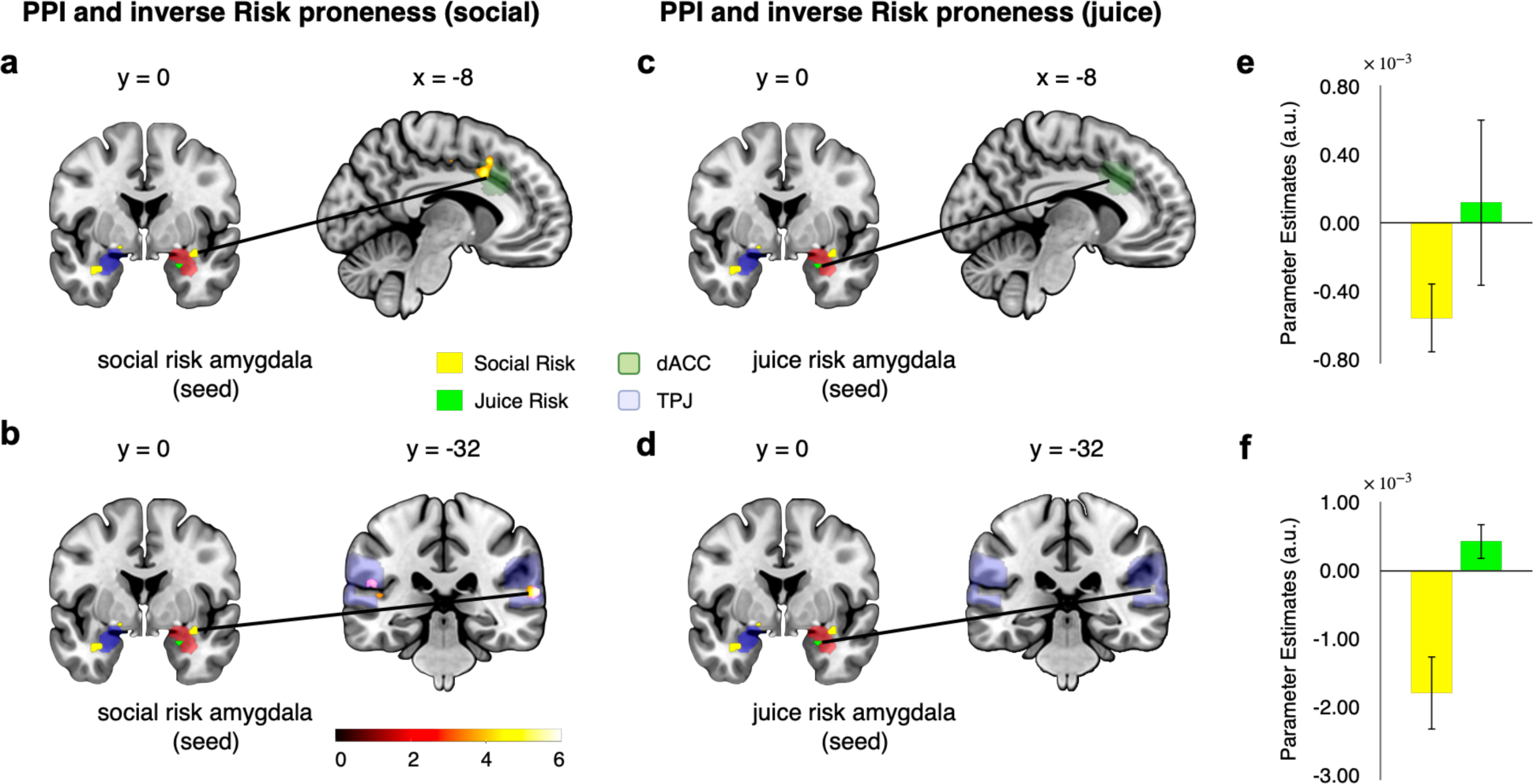
Negative correlation of amygdala-dACC connectivity with subjective risk attitude. For the PPI analysis, the region of right amygdala (yellow for social risk; green for juice risk) served as a seed. (a) The negative correlation of connectivity with social risk attitude. For social variance-risk, risk proneness correlated negatively with amygdala-dACC connectivity. Neural correlation is small volume FWE peak-level corrected p<0.05. (b) Similarly, the social risk proneness is negatively correlated to the amygdala-TPJ connectivity Neural correlation is whole-brain family-wise error (FWE) cluster-level corrected, p < 0.05, cluster-forming threshold: p<0.001. TPJ as defined by the connectivity-based bilateral anterior TPJ atlas (Mars et al., 2012). (c) In contrast, for juice risk proneness, there was no significant relations to amygdala-dACC and amygdala-TPJ connectivity (d). (e) Estimated PPI betas in dACC and (f) in TPJ were derived from both social and juice trials, with a 90% confidence interval. The parameter estimates analysis was provided from SPM12.

## 4. Discussion

We estimated the proneness of human participants to take social and non-social variance-risk using the Becker-DeGroot-Marschak auction-like procedure. We found that the amygdala processed social variance-risk independently of risk proneness. Further, the functional connectivity with social risk-coding areas dACC and TPJ during the presentation of risky choice options was negatively correlated to subjective risk proneness for social, but not for juice variance-risk.

### 4.1. Subjective risk attitude measured by willingness to pay for risky options

Although explicit valuations of options as measured with willingness to pay do not always predict choices (Abdellaoui et al., 2007), there typically is a relationship with choice, even in less consequential rating tasks (Lopez-Persem et al., 2017). Because willingness to pay is not derived from choices, it avoids any potential circularity of defining risk attitudes through choice. It is therefore noteworthy that in line with previous choice-based research (Holper et al., 2014), our participants were predominantly risk averse for non-social rewards. We extend these previous findings by showing that risk aversion is more common than risk proneness also for social rewards. Moreover, although the risk attitudes of our participants varied in both social and non-social domains, we found significant correlation between these variations across domains. These findings converge with those factor analytic approaches (Frey et al., 2017) that identified a common factor explaining general, domain-independent risk-taking tendencies. Note though that our correlations were not well-powered and should therefore be interpreted with caution.

Our approach suggests that not only the intrinsic value of social reward but also the value of the risk associated with such rewards can be measured with precise methods from behavioral economics. The intrinsic value of social reward appears to be reduced (and the amygdala response to social situations increased) in individuals with social anxiety (Schultz et al., 2019). An open question worthy of future research may be whether these individuals also show increased aversion to social variance-risk as measured by willingness to pay methods (and/or stronger amygdala responses to social risk).

From a theoretical perspective, our approach is in the finance tradition, which captures the subjective value of a choice option as a linear combination of the moments (mean, variance, etc.) that characterize the distribution of outcomes associated with that option (Markowitz, 1991). However, it is worth noting that this is not the only theory capturing value-based decision making. Indeed, both the behavior of individuals and the activity of value-related brain regions can be captured also by other theories, such as prospect theory or expected utility theory (Grabenhorst and Rolls, 2011; Konova et al., 2020; Williams et al., 2021). More importantly, our study shows that the formal approach to risk evaluation is not limited to the non-social domain but can also be applied to the social domain.

### 4.2. Amygdala encodes social and non-social variance-risk

We found social variance-risk signals in the amygdala. Although previous studies (Chang et al., 2015; Grabenhorst et al., 2019) suggest that the amygdala plays an important role in social decision making, these studies did not investigate social variance-risk. Interestingly, in our study amygdala activity specifically correlated with objectively defined social risk but not with the subjective valuation of that risk. These findings are compatible with the notion that subcortical processes are closer to the objective inputs, while cortical regions more strongly reflect the idiosyncrasies with which individuals process these inputs (Tobler et al., 2008; Genevsky et al., 2017). Together, our findings shed new light on amygdala functions in risk processing and subcortical contributions to risky decision making.

### 4.3. Subjective risk attitudes correlate with amygdala-dorsal ACC and TPJ coupling

The social variance-risk related coupling between the amygdala and dACC decreased with individual proneness to take social risks. Parts of dACC have been associated with processing social reward and social prediction errors (Burke et al., 2010a; Apps and Ramnani, 2014; Apps et al., 2015; Lockwood et al., 2015; Sul et al., 2015; Kumaran et al., 2016; Wittmann et al., 2016; Fariña et al., 2021; Westhoff et al., 2021). Specifically, the processing of social value appears to rely on the interplay of the dACC with the amygdala (Pujara et al., 2022). Similarly, our findings indicate that the TPJ, which is associated with processing social cognition (Mars et al., 2012; Hill et al., 2019; Konovalov et al., 2021), follows the same pattern of interaction. The data suggest that this interplay is relatively specific to the assessment of social, as opposed to non-social, risks.

Furthermore, we find an increasingly negative interplay between amygdala and dACC with increasing aversion to social risk during the presentation of risky choice options. This negative coupling, which is significant parameters of WTP-driven dACC activity (peak-level FWE small-volume corrected p < 0.05, cluster-forming threshold: p<0.001; see Fig. 4b and Table 3), represents a negative functional coupling with amygdala-dACC in the context of social variance-risk. These insights build upon and refine the understanding of the amygdala-dACC dynamic, previously reported to show an increasingly positive coupling associated with aversion to non-social risk in a resting state—that is, without engaging in any task (Jung et al., 2018). Our study also identified a negative functional coupling between the amygdala-TPJ and social risk proneness. Taking the two studies together, it seems that the influence of risk attitude on the interaction between these brain regions is contingent upon the active engagement in processing risk, specifically in relation to social variance-risk.

### 4.4. Conclusions

By combining functional neuroimaging with a Becker-deGroot-Marschak auction task for social and non-social outcomes we showed that the amygdala processes social variance-risk. Moreover, social risk-related coupling of the amygdala with social risk-coding areas dACC and TPJ changes as a function of risk attitude (more negative coupling with stronger risk aversion) and this relation is relatively specific for attitude to social rather than non-social risk. Our findings extend formal approaches to risk evaluation into the social domain and pave the way for investigating social risk attitude in mental-health impairment with dysfunctional processing of social uncertainty, including social anxiety.

## 5. Acknowledgements

This work was funded by the Wellcome Trust and the Royal Society (Wellcome/Royal Society Sir Henry Dale Fellowship grants 206207/Z/17/Z and 206207/Z/17/A to F.G.; Wellcome Trust Principal Research Fellowship and Programme Grant 095495 to W. Schultz). J.-C.K. received a Doc.Mobility fellowship (P1ZHP1_184166) from the Swiss National Science Foundation. P.N.T received support from the Swiss National Science Foundation (Grants 10001C_188878, 100019_176016, and 100014_165884). We thank Alaa Al-Mohammad, Philipe Bujold and Konstantin Volkmann for helpful discussions. This research was funded in whole, or in part, by the Wellcome Trust. For the purpose of Open Access, the author has applied a CC BY public copyright license to any Author Accepted Manuscript version arising from this submission.

